# A multi-enhancer *RET* regulatory code is disrupted in Hirschsprung disease

**DOI:** 10.1101/2020.06.18.159459

**Authors:** Sumantra Chatterjee, Kameko M Karasaki, Lauren E Fries, Ashish Kapoor, Aravinda Chakravarti

**Affiliations:** Center for Human Genetics and Genomics, New York University Grossman School of Medicine, New York, NY 10016; Department of Biology, Johns Hopkins University, Baltimore, MD 21205, USA; Institute of Molecular Medicine, McGovern Medical School, University of Texas Health Science Center at Houston, Houston, TX 77030, USA

## Abstract

The major genetic risk factors for Hirschsprung disease (HSCR) are three common polymorphisms within *cis* regulatory elements (CREs) of the *RET* receptor tyrosine kinase gene that reduce its gene expression during enteric nervous system (ENS) development. These variants have synergistic effects on *RET* gene expression and additionally dysregulate other ENS and HSCR genes in the *RET-EDNRB* gene regulatory network (GRN). Here, we use siRNA, ChIP and CRISPR/Cas9 deletion analyses in the SK-N-SH cell line to ask, how many HSCR-associated risk variants reside in CREs and affect *RET* gene expression? We demonstrate that 31 HSCR-associated variants reside in candidate *RET* CREs, ten with differential allele-specific *in vitro* enhancer activity and seven affecting *RET* gene expression; of these, five bind the transcription factors PAX3, RARB, GATA2 and SOX10. These and our prior results demonstrate that common sequence variants in at least 10 *RET* enhancers affect HSCR risk, extending the known *RET-EDNRB* GRN to reveal an extensive regulatory code modulating disease risk even at a single gene.

## Introduction

It is now well established that most human complex traits and diseases arise from the additive genetic effects of hundreds to thousands of variants distributed across the genome^1^. At each locus, multiple statistically significant variants are detected but it is unknown how many of them make *functionally independent* contributions to the phenotype. The widespread existence of genetic association (linkage disequilibrium, LD) between local sequence variants makes this a difficult question to answer by statistical methods alone, and requires experimental perturbation and assessment of each candidate variant ^2^. This is because genetic associations between variants depend on their recombination frequency, not their functional effects, thereby associating causal with innocent variants.

The majority of causal variants that contribute to trait variation reside within *cis* regulatory elements (CRE) and enhancers of a target gene thereby modulating its gene expression, usually in a cell-type specific manner ^3,4^. Such gene expression control is assumed to occur within a topologically associating domain (TAD) ^5,6^, defining the physical locus within which CREs function. However, three major questions remain unanswered. First, since a TAD usually harbors multiple CREs and genes, which CREs affect which gene’s expression? Second, do the different CREs of a specific gene have unique functions in space, time and cellular states or are they redundant (shadow enhancers)? Do they act independently, or are they synergistic and require clustering (super enhancers) for function ^3,7,8^? Third, since most CRE effects are small how do such small gene expression effects modulate phenotypes? The existence of many experimental methods to identify CREs comprehensively now allows us to address these questions ^9,10^.

In this study, we use Hirschsprung disease (HSCR; congenital colonic aganglionosis) as an exemplar to identify the number and activity of all ENS enhancers with disease-associated variants at its major gene *RET*. HSCR is a complex neurodevelopmental disorder in which failure of differentiation of enteric neural crest cell (ENCC) precursors during ENS development leads to aganglionosis; >35 genes/loci explaining 67% of its population attributable risk have been identified ^11^. Significantly, most of this risk arises from coding and enhancer variants of the *RET* receptor tyrosine kinase gene with smaller contributions from other genes, all of whose functions in ENS development are united through a gene regulatory network (GRN) that co-regulates *RET* and *EDNRB* ^2,11-13^. Thus, we also examine whether the CRE sequence variants we identify are individually sufficient to perturb *RET* gene expression as well as GRN activity.

Specifically, using the human neuroblastoma cell line SK-N-SH, we test 31 HSCR-associated common (minor allele frequency, MAF≥10%) polymorphisms at the *RET* locus ^14^ to demonstrate that 22 lie within CREs and of which 7 have sequence variants with differential enhancer and *RET* gene expression activity; in addition, 3 were identified before ^2^. Two of the new CREs bind the transcription factor (TF) PAX3, known to affect neural crest cell migration and ENS differentiation ^15,16^. Reducing *PAX3* expression decreases *RET* gene expression as well as that of *SOX10*, the major TF in the *RET-EDNRB* GRN ^2,17,18^. Deleting individual *RET* CREs using CRISPR/Cas9 genome editing reduces *RET* expression but is insufficient to perturb genes expression for other GRN genes. We know that *RET-EDNRB* GRN effects are evident only when *RET* gene expression falls below 50% of its wildtype level ^12^. Therefore, in HSCR, significant reductions of gene expression at *RET* and its GRN to affect ENS development only occurs in individuals with multiple CRE variants in combination with variants in other GRN genes. The corollary is that a diverse regulatory code affects complex disease genes.

## Results

### Enhancers at the RET locus

To create a complete catalog of common (MAF≥10%) *RET* regulatory variants associated with HSCR, we began with analysis of all 38 genome-wide significant non-coding single nucleotide polymorphisms (SNPs) discovered in a genome-wide association study (GWAS) of 220 HSCR trios comprising a proband and both of her/his parents ^14^. These SNPs were distributed across 6 LD blocks in a 155kb TAD (chr10: 43434933-43590368; hg19) containing *RET* as the sole gene (**Figure 1A**). We previously analyzed 8 of these SNP-containing genomic elements because they each disrupted a predicted TF binding site (TFBS) determined form ENCODE ChIP-seq data ^2^.

**Figure 1:**
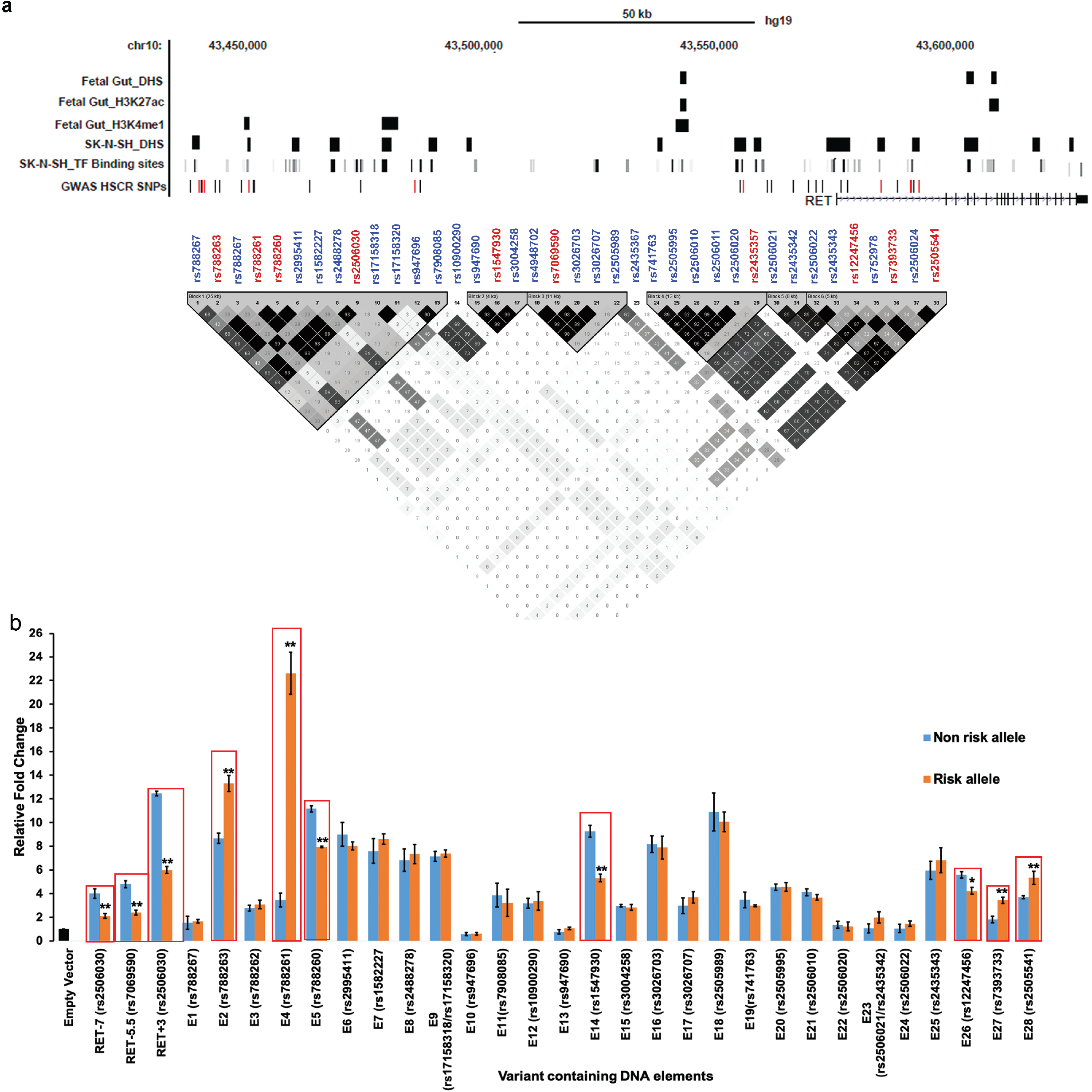
The *RET* regulatory landscape in the enteric nervous system. (A) The 155kb *RET* locus (10:43434933-43590368; hg19) contains 38 putative cis regulatory elements (CRE) with HSCR-associated polymorphisms in 4 linkage disequilibrium (LD) blocks. Multiple enhancer-associated epigenetic marks (DHS: DNaseI hypersensitivity, H3K27ac, H3K4me1) in 108-day human fetal large intestine and the SK-N-SH neuroblastoma cell line, and transcription factor (TF) binding sites, from public sources, are noted. All common (≥10% allele frequency) variants associated with HSCR are shown, with those showing allelic difference in *in vitro* transcription assays marked in red (this paper). (B) Allele specific *in vitro* luciferase assays of the 38 CREs in (A) in SK-N-SH cells are shown: 22 CREs act as enhancers, as compared to a promoter-only control, of which, 7 also show allelic difference in luciferase activity (boxed) for its cognate HSCR-associated polymorphism in addition to RET-7, RET-5.5 and RET+3 positive controls. Error bars are standard errors (SE) of the mean (*P < 0.01, **P < 0.001) for 3 independent biological replicates for each assay.

We first asked: do the 30 remaining SNPs resided within CREs? We conducted *in vitro* functional tests of enhancer activity by cloning ∼500 bp elements centered on each risk variant (**Table 1**) into a pGL4.23 luciferase vector, with a minimal TATA-box of the β-globin gene and transfecting them into the human neuroblastoma SK-N-SH cell line. SK-N-SH expresses all known members of the *RET-EDNRB* GRN and is an appropriate cell model system for studies of ENS transcriptional regulation ^2,12^. Two pairs of SNPs (rs17158318/ rs17158320 and rs2506021/ rs2435342) were only 64bp and 108bp apart and were cloned into the same elements (E9 and E24, respectively) (**Table 1**). As positive controls, we re-analyzed 3 HSCR associated SNPs (rs2506030, rs7069590, rs2435357) previously shown to be *RET* enhancer variants ^2^ (**Table 1**). Our reporter assays showed that 22 new elements had significant enhancer activity (*P* <0.001 and >2X reporter activity over the promoter only control vector) of which 7 (E2, E4, E5, E14, E26, E27 and E28) also displayed differential reporter activity between the risk and non-risk alleles (**Table 1 and Figure 1B**). Among the latter, 71% (E2, E4, E26, E27 and E28) overlapped an open chromatin region or an enhancer-associated epigenetic mark in the human fetal gut and SK-N-SH ^19,20^ while 20% (E9, E16, E21) of the remaining 15 elements with reporter activity that had no allelic difference overlapped a potential enhancer mark (**Figure 1A**). Thus, at least 10 functionally distinct CREs within the *RET* TAD can potentially affect HSCR risk.

**Table 1:**
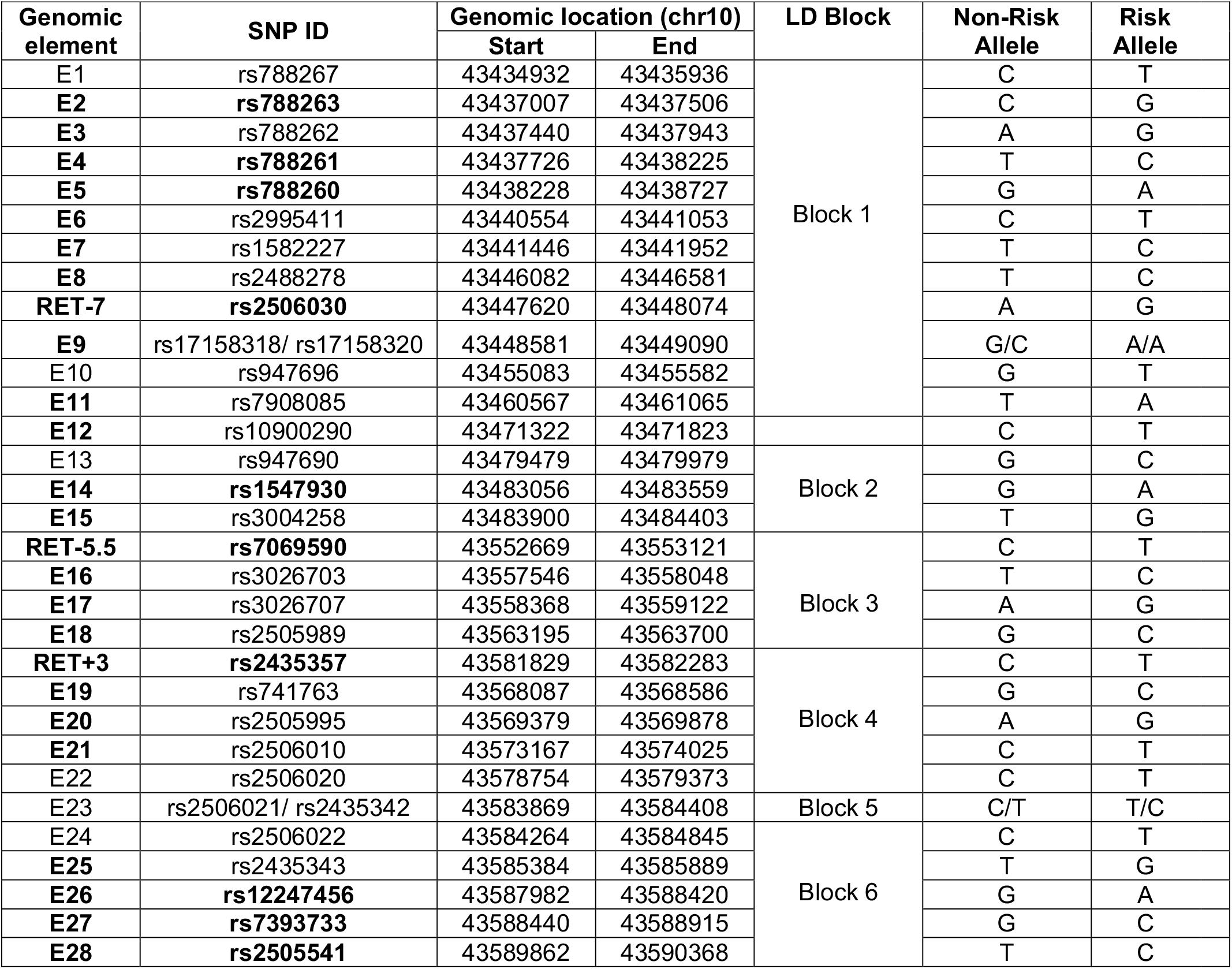
Genomic coordinates (hg19) of 31 elements containing 33 single nucleotide polymorphisms (SNPs), within 6 linkage disequilibrium (LD) blocks at the *RET* locus which are associated with Hirschsprung disease (HSCR). 25 elements in bold act as enhancers in *in vitro* transcriptional assays while 10 SNPs in bold show allelic difference between the risk and non-risk allele as well. Enhancer activities of elements RET-7 (rs2506030), RET-5.5 (rs7069590) and RET+3 (rs2435357) have been previously demonstrated to be affected by the HSCR associated risk alleles^2^.

### Haplotype specific effect of causal RET polymorphisms

The identification of 10 CRE-associated risk variants (rs788263, rs788261, rs788260, rs2506030, rs1547930, rs7069590, rs2435357, rs12247456, rs7393733 and rs2505541; **Table 1**) prompted us to ask which allelic combinations were associated with disease risk. First, we estimated haplotypes and their frequencies for all 10 SNPs in 220 unrelated HSCR cases ^14^ and 503 unrelated controls from the 1000 Genomes project ^21^, all of non-Finnish European-ancestry. Second, we estimated the odds ratio (OR) for all haplotypes with a frequency ≥1% in controls. We observed 10 distinct haplotypes of which CTGAACCAC**T** (risk allele in bold) was used as the reference because it had the smallest number (one) of risk alleles (**Table 2**) and we have previously shown that HSCR risk scales with increasing number of CRE variants ^2,11-13^: significant risk was observed for two haplotypes, **GCAGGTTGGT** (OR 12.2, 95% CI: 5.97-24.93, P=7.02×10^−12^) and CTGA**GTTGGT** (OR 7.2, 95% CI: 3.26-15.91, P=1.02×10^−6^) (**Table 2**). Unsurprisingly, these two haplotypes contain our previously identified risk increasing ATT and GTT haplotypes (for rs2506030, rs7069590, rs2435357) ^2^. The 10-SNP risk haplotypes differ only for the first four SNPs (rs788263, rs788261, rs788260, rs2506030) which occur within the most 5’ LD block. SNPs within this LD block do contribute to HSCR but we do not have a sufficient sample size to test our hypothesis that **GCAGGTTGGT** (OR 12.2) has a significantly higher risk than CTGA**GTTGGT** (OR 7.2). Even discounting these variants, HSCR risk is clearly spread over at least three LD blocks suggesting multiple independent enhancer variants contributing to risk (**Figure 1**). This is *prima facie* evidence that these SNPs contribute to risk synergistically. In other words, risk of or protection from HSCR depends on the simultaneous binding or lack-of-binding of multiple independent TFs at *RET* CREs, implying a *RET* regulatory code.

**Table 2:** Haplotypes for ten *RET* enhancer polymorphisms (rs788263, rs788261, rs788260, rs2506030, rs1547930, rs7069590, rs2435357, rs12247456, rs7393733 and rs2505541), with risk alleles denoted in bold. Observed frequencies of haplotypes among 220 European ancestry HSCR cases and 503 controls together with the odds ratio (with respect to the reference haplotype CTGAACCACT containing only one susceptibility allele; significant values bolded) and statistical significance (P) are shown.

| Haplotype | Case Frequency | Control Frequency | Odds Ratio (95% CI) | P |
| --- | --- | --- | --- | --- |
| CTGAACCACT | 0.02 | 0.08 | 1 | 1 |
| CTGAATCACT | 0.04 | 0.07 | 2.24 (0.95-5.29) | 0.07 |
| CTGAGCCACT | 0.03 | 0.10 | 1.13 (0.45-2.82) | 0.79 |
| CTGAATCGGC | 0.04 | 0.06 | 2.84 (1.20-6.77) | 0.02 |
| CTGAGTCGGC | 0.06 | 0.16 | 1.58 (0.71-3.51) | 0.26 |
| CTGAATTGGT | 0.02 | 0.01 | 4.35 (1.41-13.49) | 0.01 |
| CTGAGTTGGT | 0.10 | 0.06 | **7.2 (3.26-15.91)** | **1.02x10<sup>-6</sup>** |
| **GCAGGTC**ACT | 0.02 | 0.04 | 2.45 (0.92-6.54) | 0.07 |
| **GCAGGTC**GGC | 0.06 | 0.15 | 1.62 (0.72-3.61) | 0.24 |
| **GCAGGTT**GGT | 0.52 | 0.17 | **12.20 (5.97-24.93)** | **7.02x10<sup>-12</sup>** |

### Transcription factors regulating RET

To identify the TFs that underlie this code, we searched for TFBSs within the 7-novel risk-associated CREs and identified four candidate TFs: (1) PAX3 binding to E2 (AATAAACCC; P=4.67×10^−5^) and E27 (TCGTCACTCTTAC; P= 9.99×10^−5^), (2) ZBTB6 to E5 (TGGCTCCATCATG; P=2.387×10^−6^), and, (3) ZNF263 binding to E14 (GCCTCACTGCTCCAG; P=8.09×10^−5^). To determine their relevance for HSCR, we performed qPCR in SK-N-SH cells and observed no expression for *ZNF263* and *ZBTB6* (data not shown). Further, their expression was absent in the developing mouse gut ^22^, where *RET* expression is critical for ENS development ^23^, making it highly unlikely that these TFs control *RET* expression via specific enhancers. Moreover, *ZNF263* and *ZBTB6* are both zinc finger domain-containing proteins that are GC-rich, a feature which overestimates their statistical significance owing to their rarity.

In contrast, we detected robust expression of *PAX3* in both SK-N-SH and the developing mouse gut ^22^. Thus, we performed ChIP-qPCR for *PAX3* in SK-N-SH cells and detected significant binding at both E2 (18-fold enrichment; P= 10^−3^) and at E27 (26-fold enrichment, P= 5×10^−4^) (**Figure 2**). We further demonstrated the specificity of this binding by performing ChIP-qPCR after siRNA-mediated knockdown of PAX3 in SK-N-SH cells to show a 1.3-fold (P=8×10^−3^) reduced binding at E2 and 2-fold (P=4×10^−4^) reduced binding at E27 (**Figure 2**). To further prove that PAX3 does indeed control *RET*, we also measured *RET* gene expression after siRNA-mediated knockdown of *PAX3*. As positive controls, we measured *RET* levels after siRNA-mediated knockdown of the established *RET* TFs SOX10, GATA2 and RARB. These experiments showed that decreasing *PAX3* led to a 49% (p=4×10^−4^) reduction in *RET* in comparison to 76% (p=2.3×10^−6^), 50% (p= 3.1×10^−3^) and 81% (p= 4.1×10^−5^) decreases consequent to *SOX10, GATA2* and *RARB* knockdown, respectively; as a control, knockdown of *RET* by its specific siRNA reduced its expression by 96% (p=4.4×10^−6^) (**Figure 3A**). We have previously demonstrated that there is considerable cross-talk between the established RET TFs ^2^. Consequently, we measured gene expression of *SOX10, GATA2* and *RARB* after siRNA-mediated knockdown of *PAX3*: we observed only a significant drop in *SOX10* gene expression (32% decrease, P= 3×10^−3^); *GATA2* and *RARB* levels were decreased but not significantly so (**Figure 3B)**.

**Figure 2:**
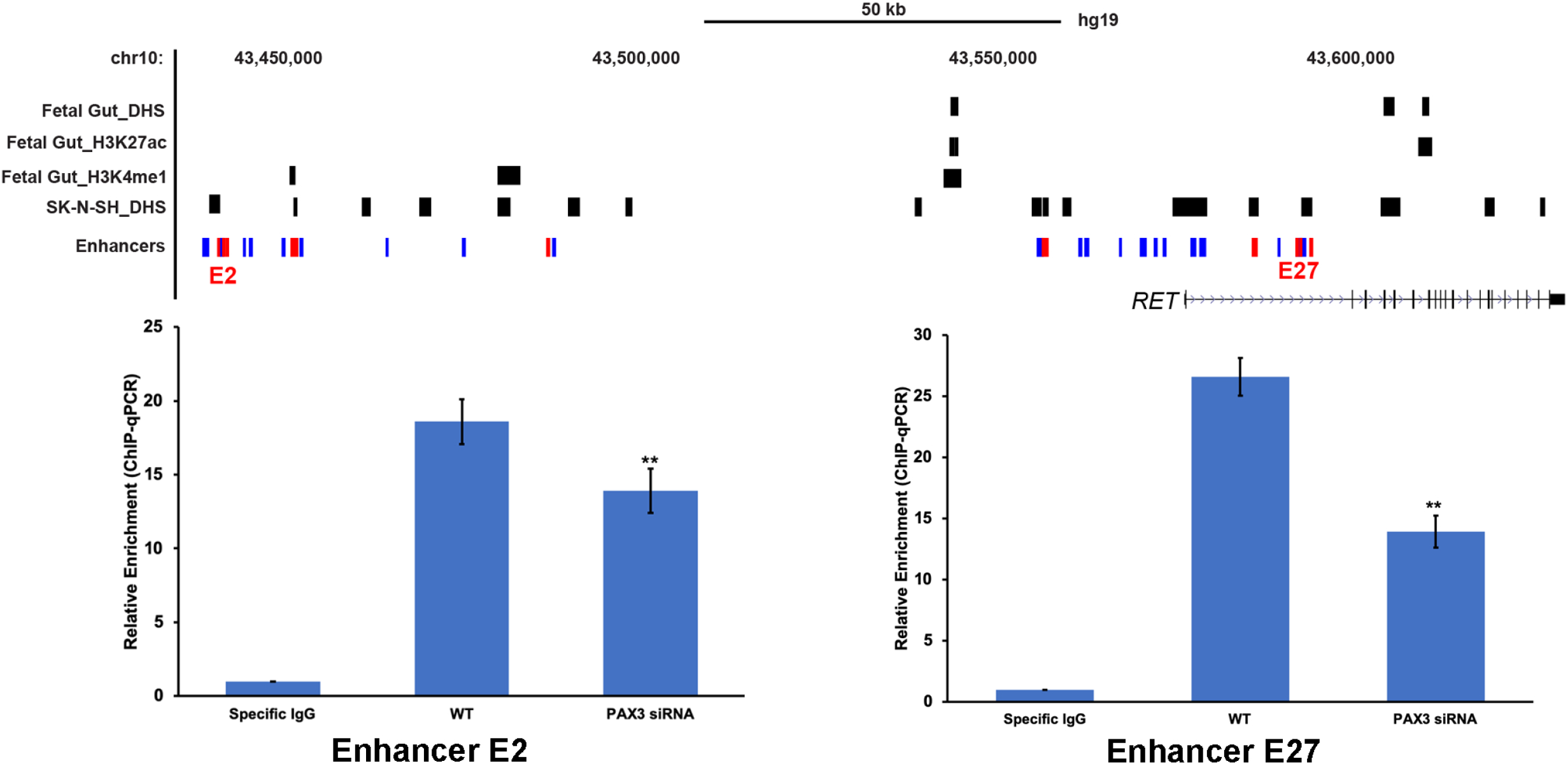
Identification of cognate transcription factors bound to *RET* enhancers. Genome map of the *RET* locus with locations of the E2 and E27 CREs together with ChIP-qPCR results using a PAX3 antibody in SK-N-SH cells shows enrichment of binding as compared to the background. The specificity of binding is shown by siRNA knockdown of *PAX3* with concomitant reduction in ChIP-qPCR signals at both CREs. Error bars are standard errors (SE) of the mean (*P < 0.01, **P < 0.001) for 3 independent biological replicates for each ChIP assay.

**Figure 3:**
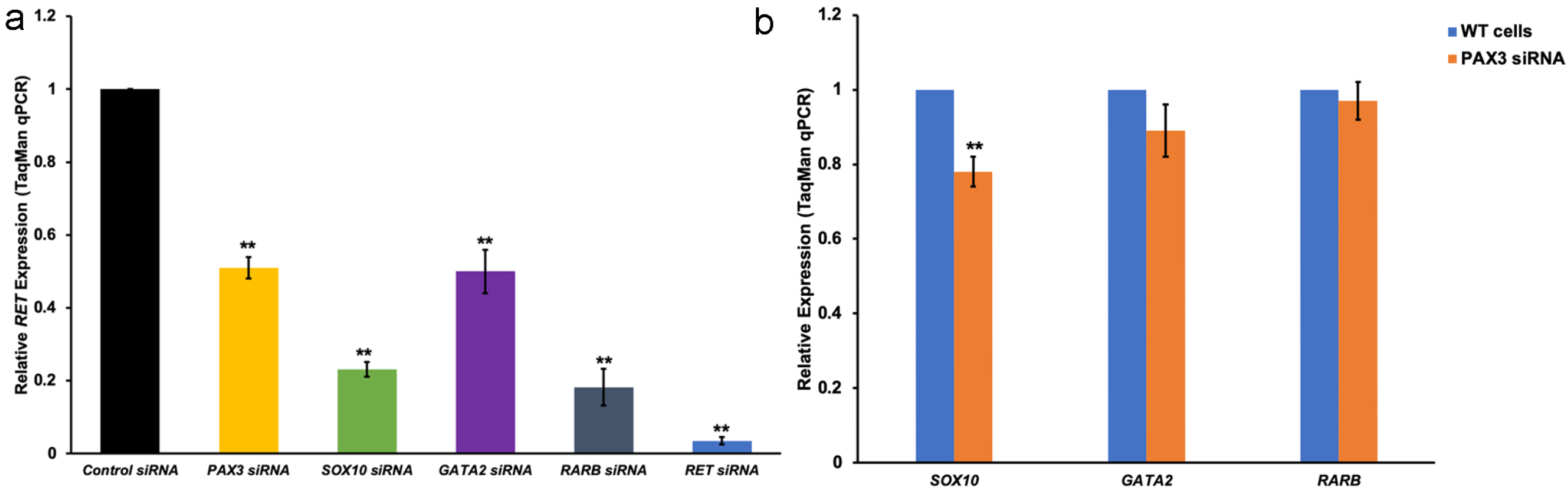
Transcription factor-mediated *in vitro* and *in vivo* effects on gene expression. (A) siRNA-mediated knockdown of *PAX3, SOX10, GATA2, RARB and RET* in SK-N-SH cells decreases *RET* gene expression. (B) siRNA-mediated knockdown of *PAX3* has significant transcriptional effects on *SOX10* but small yet statistically insignificant effects on *GATA2* and *RARB*. Error bars are standard errors (SE) of the mean (*P < 0.01, **P < 0.001) for 5 independent biological replicates in all experiments.

### in vivo evidence for RET enhancers

The human genetic evidence for HSCR-associated polymorphisms within CREs identified from *in vitro* (reporter activity) and *ex vivo* (siRNA in SK-N-SH) experiments can be buttressed by deletion analysis of each enhancer rather than knockdown of its cognate TF. To do so, we designed a single guide RNA close to each HSCR-associated SNP for all 10 target enhancers to introduce non-homologous end joining-induced deletions in SK-N-SH cells. We screened 5 independently transfected wells for each guide and verified them by Sanger sequencing: none of the deletions were > 10bp. Except for enhancer E5, successful deletions were observed in the remaining CREs encompassing the polymorphic site. We used the *Inference of CRISPR Edits (ICE)* tool ^24^ to estimate that individual guides introduced deletions >3bp in 10-50% of the cells in all successfully targeted CREs (**Supplementary Table 1**).

We next measured *RET* gene expression in these enhancer-deleted cells. Our results demonstrate that except for enhancers E2 and E14, deletion of DNA sequences surrounding the HSCR-associated SNPs in all other CREs led to changes in *RET* expression. Thus, deletion of E4 (24%; p=3.2×10^−4^) leads to higher expression while deletion of E26 (28%; p= 3.7×10^−4^), E27 (19%; p=1.2×10^−3^) and E28 (29%; p=3.2×10^−4^) all lead to lower *RET* expression (**Figure 4A**). The positive controls, RET-7 (22%; p=1.3×10^−3^), RET-5.5 (22%; p=2×10^−3^), and RET+3 (32%; p=4.1×10^−4^) reduced *RET* gene expression as expected. It is interesting that all 4 intronic enhancers (RET+3, E26, E27 and E28) residing within a 8.5 kb region (chr10:43,581,812-43,590,347) in the first intron of *RET* independently control *RET* expression. Thus, these elements might be a part of a single *enhanceosome* critical for spatio-temporal expression of *RET*; note that the risk-associated variants at these sites are on a single haplotype (**Figure 1A**).

**Figure 4:**
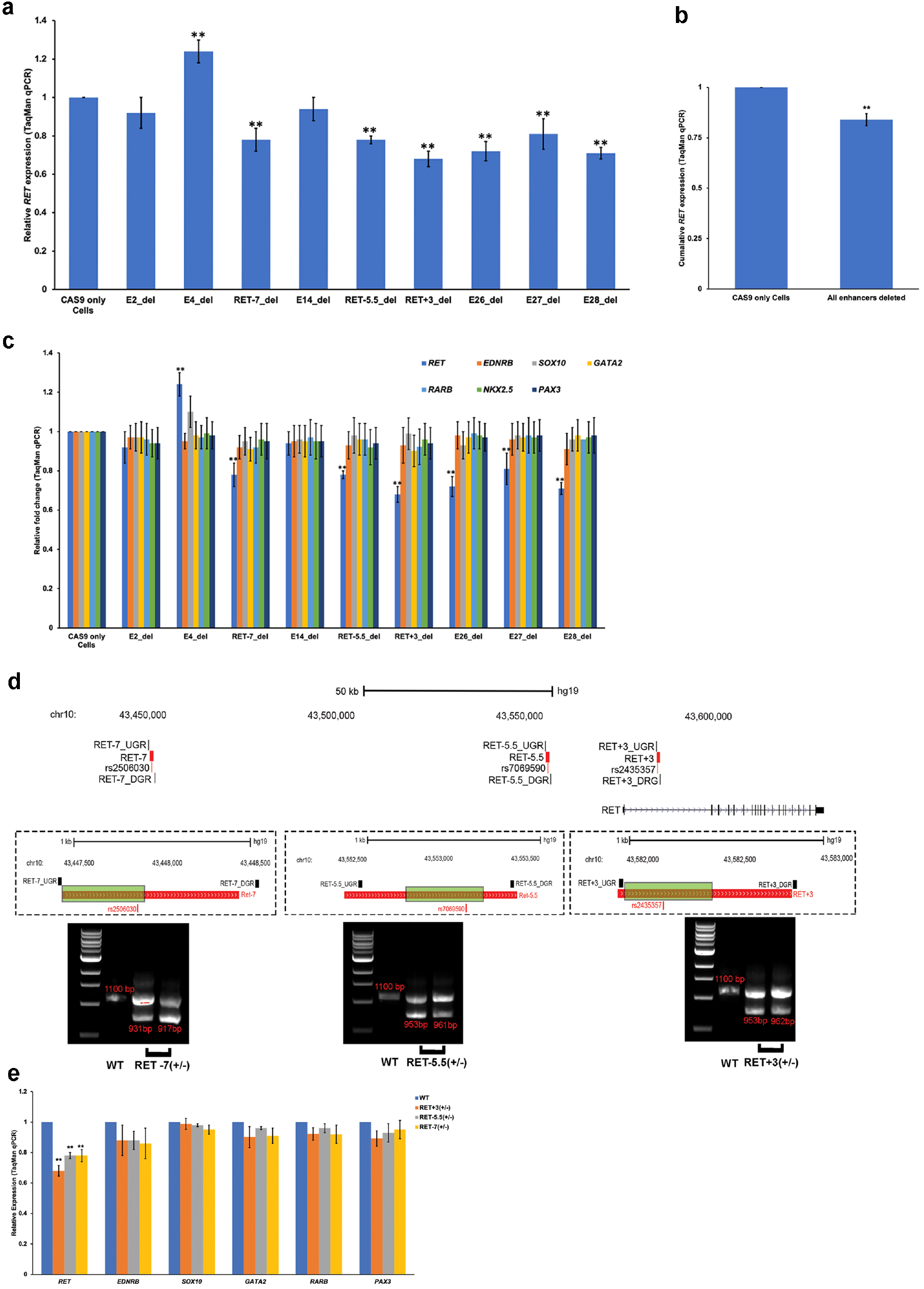
CRISPR/Cas9 induced deletions of *RET* enhancers with HSCR-associated variants reduce *RET* gene expression *in vitro*. (A) There is significant loss of *RET* gene expression from 7 of 9 CREs with small (≤10bp) deletions centered on the variant site; only the E14 enhancer shows increased gene expression. (B) There is significant loss of average *RET* expression from the cumulative effect of individual enhancer deletions as compared to wildtype cells. (C) The expression of other *RET* GRN genes are unaffected by these CRE deletions likely owing to *RET* expression never decreasing below 50%. (D) A genomic map of the *RET* locus showing 3 positive control CREs and their corresponding HSCR-associated polymorphisms: RET-7 (rs2506030), RET-5.5 (rs7069590) and RET+3 (rs2435357). The green boxed region is the deleted region for each enhancer as confirmed by Sanger sequencing and validated by agarose gel electrophoresis. (E) Heterozygous deletion of all 3 enhancers lead to significant loss of *RET* gene expression but not other *RET* GRN genes. Error bars are standard errors (SE) of the mean (*P < 0.01, **P < 0.001) for 5 independent biological replicates in all experiments for single guide deletions, across 6 replicates for RET-7 and 9 replicates for RET-5.5 and RET+3 in dual guide deletions.

HSCR is associated with loss of *RET* function ^25,26^ and diminished *RET* expression leads to loss of ENS during gut development in mice, the hallmark of HSCR ^22,27^. Thus, we predict that the cumulative effect of the disruption of all HSCR-associated CRE at E4, RET-7, RET-5.5, RET+3, E26, E27 and E28 should lead to reduced *RET* expression. Normalized *RET* expression averaged over individual enhancer deletion experiments indeed demonstrates that *RET* expression is diminished by 16% (p= 0.003) when compared to CAS9 only transfected (wildtype) cells (**Figure 4B**). Thus, irrespective of the direction of change in *RET* expression due to individual deletion, the cumulative disruption of these enhancers leads to *RET* deficiency.

We next asked whether these changes in *RET* expression had concomitant changes in the expression of the remaining members of the *RET* GRN by quantifying expression of *EDNRB, SOX10, GATA2, RARB, NKX2*.*5 and PAX3*, the members of the GRN which expresses in this cell line, in the individual enhancer-deleted cells. Our results demonstrate no significant changes in gene expression (**Figure 4C**). This result is not unexpected given our previous finding that GRN transcriptional dysregulation occurs only when *RET* gene expression falls below 50% of wildtype levels ^2,12^.

Since the individually strongest HSCR associations were within the previously discovered RET-7, RET-5.5 and RET+3 enhancers (OR of 1.8, 1.7 and 4.1 respectively) ^2^, we wanted to ascertain if DNA sequences beyond the polymorphic site impacted *RET* expression. Thus, we generated larger deletions in these enhancers using paired guide RNAs to direct espCAS9 to these specific DNA sequences in SK-N-SH (**Figure 4D**). We screened 48 independently transfected wells for each enhancer and detected only heterozygous deletions for each. For RET-7, we identified 2 clones (4%) with 169bp and 183bp deletions, both of which deleted rs2506030. We detected 3 heterozygous clones for RET-5.5 deletion (6.25%): 2 clones with 147bp and 1 clone with a 139bp, all of which deleted rs7069590. Similarly, we also identified 3 clones (6.25%) with RET+3 deletion with 1 clone encompassing 147bp and 2 clones encompassing 138bp deletions, all of which deleted rs2435357(**Figure 4D**). Next, we measured gene expression of *RET, EDNRB, SOX10, GATA2, RARB, NKX2*.*5 and PAX3* by qPCR in these cells to detect the effect of heterozygous enhancer deletions on the expressed genes of the *RET* GRN. Deletion of RET+3 led to 32% (P=3.1×10^−3^) loss of *RET* expression compared to cells transfected with only a Cas9 vector without the guide RNA; the corresponding decreases for RET-5.5 and RET-7 were 22% (P=2.8×10^−3^) and 20% (P=2.6×10^−3^), respectively, consistent with their individual risks (OR of 4.1, 1.7 and 1.8, respectively) (**Figure 4E**) (Chatterjee et al., 2016; Emison et al., 2010; Kapoor et al., 2015). Thus, deletion of sequences beyond the region immediately surrounding the polymorphisms do not seem to have any substantial effect on *RET* expression. Thus, for HSCR, these SNPs are the causal risk factors. Here as well, the remaining genes of the GRN are transcriptionally unchanged (**Figure 4E**) because *RET* expression never falls below 50%.

## Discussion

It is evident that a multiplicity of enhancers control a gene’s expression: this feature has many implications for complex disease genetics and its mechanisms. The data reported here, based on human genetics, siRNA, and ChIP analyses in the SK-N-SH cell line, together with our prior studies ^2,11-13^, have identified 38 distinct CREs of *RET* with sequence variants that are associated with HSCR. Of these, 10 contain common polymorphisms with differential allelic enhancer activity. CRISPR/Cas9 deletion analyses of these CREs demonstrate that at least 7 of these *RET* enhancers, binding the SOX10, RARB, GATA2, PAX3 and yet unknown TFs, have all of the hallmarks of harboring causal variants for HSCR (**Table 3**). Given that none of these experimental approaches are 100% efficient yet other enhancers are likely. Nevertheless, the many enhancers at the *RET* locus, identified through *in vitro* analyses, do not imply that all of them are involved in gut (as opposed to other tissue) development, and, if so, whether they have unique roles in the ENS across development, as we have shown earlier (Chatterjee et al., 2016) or are merely shadow enhancers ^8^. *RET* is highly expressed in the mid- and fore-brain, kidney and dorsal root ganglia: thus, these CREs may regulate *RET* in these tissues.

**Table 3:**
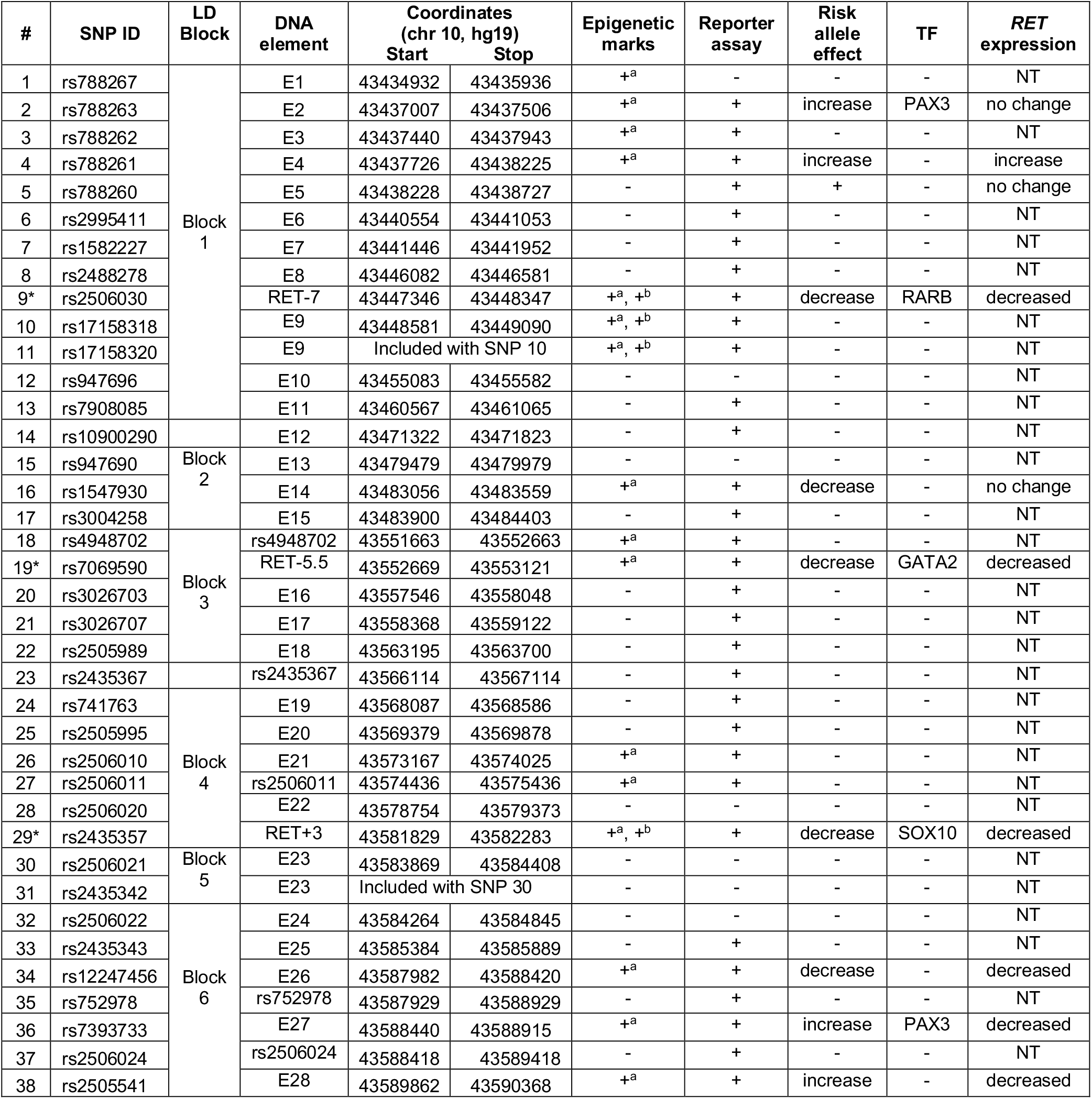
38 Hirschsprung disease (HSCR)-associated polymorphisms in 6 linkage disequilibrium (LD) blocks, contained within 36 DNA elements at the *RET* locus annotated with respect to epigenetic marks (DNaseI hypersensitivity (+^a^) in the SK-N-SH cell line, H3K4me1 (+^b^) marks in human fetal gut), luciferase reporter assays of alleles in the SK-N-SH cell line, allelic differences in luciferase assays in the SK-N-SH cell line, the transcription factor (TF) binding the indicated regulatory element, and whether deletion of the element affected *RET* gene expression. 31 elements have enhancer activity in *in vitro* luciferase assays, 10 of which demonstrated affect due to the HSCR associated risk polymorphisms. CRISPR based *in vivo* deletion of these 10 enhancers identified 7 which affect *RET* gene expression, 5 of which are bound by PAX3, RARB, GATA2 and SOX10 TFs. Elements #9, 18, 19, 23, 27, 29, 35 and 37 were reported in our previous study^2^ and are included for completeness; of these 9,19 and 29, marked by *, are positive controls with *in vivo* evidence of enhancer activity in transgenic mice. NT: not tested.

Definitive proof of our hypothesis that the 7 *RET* enhancers cause HSCR will require the creation of mouse models with multiple regulatory variants at the same locus, far more feasible using new methods of synthetic biology ^28^ as opposed to CRISPR/Cas9 genome editing. Nevertheless, this hypothesis is supported by the observation that RARB, GATA2, SOX10 and PAX3 binding to five of these enhancers have known roles in ENS development and HSCR ^15,16,29^. In other words, if the trans factors lead to HSCR and the cis factors that bind them are HSCR-associated then these cis factors are direct risk factors of HSCR. Their distinct nature and LD relationships also suggests that they contribute independently to *RET* gene expression, and, therefore to HSCR.

The multiplicity of noncoding variants in CREs all controlling the same gene should give us pause in interpreting the effect size and functional effects of individual GWAS variants. As we have shown, the cumulative effect of all variant-bearing enhancers is to *significantly lower RET* gene expression, as expected from *RET* coding mutations ^17,18^. As we have also demonstrated, the largest risks are associated with haplotypes with multiple risk alleles, even across LD blocks ^2^. This accumulation of multiple CRE variants is expected to lead to a larger effect through reduced binding of multiple TFs to multiple enhancers. Given the role of multiple TFs on the promoter this is likely synergistic. Additionally, other interactions are important in ENS cells with these genotypes. Note that *RET* also controls the gene expression of its own TFs PAX3, RARB, GATA2 and SOX10. This feedback may be a secondary but an important cause of reduced *RET* gene expression further exacerbating the enhancers’ effect. We do not yet know whether this diversity of genetic control with feedback is typical or not. *RET* is a highly dosage-sensitive gene with higher and lower than wildtype levels being associated with neuroendocrine tumors and aganglionosis, respectively. Although extensive regulatory control is common for developmental genes^30^, the genetic lessons are likely to be universal.

The human genetic implications of these data, beyond understanding HSCR, are that genetically independent (unassociated) SNPs at a specific GWAS locus are not the only candidate variants for understanding a phenotype. Additional SNPs may be involved, even those perfectly associated with one another provided they affect functionally independent enhancers, as has been demonstrated by the recent discovery of additional *RET* variants and CREs that control its expression ^31,32^. Finally, such regulatory control by the variant allele can also be quite varied (E4, E27 versus others), decreasing versus increasing enhancer activity, depending on activator versus repressor TFs and their co-regulators, as well as decreasing as well as increasing target gene expression. This suggests that understanding the regulatory contributions to GWAS will require experimental data on enhancer effects beyond what statistical analysis can suggest. Furthermore, we need broader enhancer screens to define the full enhancer architecture of *RET* and do so *in vivo* at different developmental stages and by sex. We also need to elucidate the full repertoire of TFs that regulate *RET*. These pieces of information are crucial to understand the full extent and composition of the *RET-EDNRB* GRN, which in turn will identify new genes that become mutational targets of HSCR.

## Materials and methods

### Cell lines

The human neuroblastoma cell SK-N-SH, purchased from ATCC, USA (# HTB-11), was grown under standard conditions (DMEM + 10% FBS and 1% Penicillin Streptomycin). It was maintained in 10 cm culture dishes and passaged every 48 hours when it reached ∼80% confluency.

### ChIP-seq peak calling

Three epigenomic datasets generated from a 108-day human fetal large intestine, histone H3K27ac ChIP-seq (GSM1058765), histone H3K4me1 ChIP-seq (GSM1058775), and DNaseI-seq (GSM817188), were downloaded from the NIH Roadmap Epigenomics Project ^19^. For the SK-N-SH cell line, DNaseI-seq data (GSM736559) were obtained from the ENCODE project ^33^. For each dataset, MACS software v1.4 ^34^ with default settings was used to call ‘‘peaks’’ at genomic sites where sequence reads were significantly enriched over background. With the default peak-calling threshold (P < 10^−5^), 51,771, 61,689, 66,930 and 52,534 genomic regions were identified in the GSM1058765, GSM1058775, GSM817188 and GSM736559 datasets, respectively.

### Reporter assays

400 ng of firefly luciferase vector (pGL4.23, Promega Corporation,USA) containing the DNA sequence of interest and 2 ng of Renilla luciferase vector (transfection control) were transiently transfected into the SK-N-SH cell line (5 – 6×10^4^ cells/well), using 6 ml of FuGENE HD transfection reagent (Roche Diagnostic, USA) in 100ml of OPTIMEM medium (Invitrogen, USA). Cells were grown for 48 hours and luminescence measured using a Dual Luciferase Reporter Assay System on a Tecan multi-detection system luminometer, per the manufacturer’s instructions.

### Chromatin Immunoprecipitation–qPCR (ChIP-qPCR) assays

ChIP was performed thrice independently for each antibody using 1×10^6^ SK-N-SH cells for each transcription factor using the EZ-Magna ChIP kit (Millipore Sigma, USA), as per the manufacturer’s instructions, with the following modifications: the chromatin was sonicated for 30s on and 30s off for 10 cycles; sheared chromatin was pre-blocked with unconjugated beads for 4 hours and specific antibodies separately conjugated to the beads for 4 hours before immunoprecipitation was performed with the pre-blocked chromatin. A polyclonal antibody was used against PAX3 (16HCLC; Invitrogen, USA) at 15μg concentration. ChIP assays were also performed on cells 48 hours after transfection with the PAX3 siRNAs (L-012399-00-0005; Dharmacon/Horizon Discovery, USA) at 25 μM to assess specificity of TF binding. qPCR assays were performed using SYBR green (Life Technologies, USA) and specific primers against enhancer E2 (E2_FWD 5’ GCTGCAGATATGCAACTTCCAA 3’ and E2_REV 5’ AGATATGCTGGTGAGGGGCT 3’) and enhancer E27 (E27_FWD 5’ AGGAAGGTAGGCACCCTGTA and E27_REV 5’ AGCCCTGTGTTAACTGTCCG 3’). The data were normalized to input DNA and enrichment was calculated by fold excess over ChIP performed with specific IgG as background signal. All assays were done in triplicate for each independent ChIP assay (n = 9).

### siRNA assays

*PAX3* (L-012399-00-0005), *RET* (L-003170-00-0005), *SOX10* (L-017192-00), *GATA2* (L-009024-02 and *RARB* (L-003438-02) SMARTpool siRNAs (combination of 4 distinct siRNAs targeting each gene) along with ON-TARGET plus non-targeting siRNAs (D-001810-10, negative control) (Dharmacon/ Horizon Discovery, USA) were transfected at 20 μM in SK-N-SH cells at a density of 10^4^ -10^5^ cells using FuGene HD Transfection reagent (Promega Corporation, USA) per the manufacturer’s instructions. Negative control siRNAs were always transfected at 25 μM concentration.

### Gene expression assays

Total RNA was extracted from SK-N-SH cells using TRIzol (Life Technologies, USA) and cleaned on RNeasy columns (QIAGEN, USA). 500μg of total RNA was converted to cDNA using SuperScriptIII reverse transcriptase (Life Technologies, USA) using Oligo-dT primers. The diluted (1/5) total cDNA was subjected to Taqman gene expression (ThermoFisher Scientific, USA) using the following transcript-specific probes and primers: *RET* (Hs01120032_m1), *EDNRB* (Hs00240747_m1), *PAX3*(Hs00240950_m1), *SOX10* (Hs00366918_m1), *GATA2* (Hs00231119_m1) and *RARB* (Hs00977140_m1)

Human β-actin was used as an internal loading control for normalization. For siRNA knockdown experiments, five independent wells of SK-N-SH cells were used for RNA extraction and each assay performed in triplicate (n = 15). Relative fold change was calculated based on the 2ΔΔCt (threshold cycle) method. For siRNA experiments, 2ΔΔCt for negative control non-targeting control siRNA was set to unity. P values were calculated from pairwise 2-tailed t tests and the data presented as means with their standard errors (SE).

### CRIPSR/Cas9 induced deletions

#### (i) Single guide disruptions

Each enhancer region centered on a polymorphic site was targeted using a single guide RNA (**Supplementary Table 2**) by transfecting a ribonucleoprotein complex containing 100pmol of specific gRNA, coupled with 5μg/μl of Truecut cas9 nuclease (ThermoFisher Scientific) in Lipofectamine Crisprmax solution (ThermoFisher Scientific). For each enhancer, 5 wells containing ∼30,000 SK-N-SH cells were independently transfected. To increase efficiency of deletion, we re-transfected the cells with the same ribonucleoprotein mix a second time after 72 hours. The cells were further grown for 48 hours and then equally split into 2 tubes for DNA and RNA extraction. To confirm disruption of the enhancer regions, specific primers (**Supplementary Table 3**) were used to verify deletions by PCR followed by Sanger sequencing. We used the *Inference of CRISPR Edits (ICE)* tool ^24^ to estimate the percentage of cells carrying various insertion/deletions.

#### (ii) paired guide disruptions

The following genomic regions were targeted with pairs of guides for 3 enhancers: RET-7 (*RET-7_DGR-5’GACCCCTCTGGACCCGTCGCGG3* and *RET-2_UGR-CCGAGTGTTTGCGCCCATGAGG*), RET-5.5 (*RET-5*.*5_DGR-AAGGATGGATACACCTTCCGGG* and *RET-5*.*5_UGR-AAGGATGGATACACCTTCCGGG*) and RET+3 (*RET+3_DGR-GCCTGGCTGGAGGTCCAAGAGG* and *RET+3_UGR-GCTCATGAGGAGCACACCGTGG*). The guide RNA and Cas9 were cloned into the vector *U6-gRNA/CMV-Cas9-GFP* (Millipore Sigma, USA) and each pair of guides were transfected at 10μg into 72 wells of low-density (∼10,000 cells) plated SK-N-SH cells. We also transfected 72 wells with only the Cas9-GFP vector without the guide RNAs to serve as controls against which to measure gene expression changes arising from our enhancer deletions.

After 24 hours cells were checked for GFP expression and wells with at least ∼80% positive cells were re-plated, ensuring high transfection efficiency. For all 3 constructs and the control we obtained at least 48 wells with over 80% transfection efficiency. The cells were further grown for 24 hours and then split into 2 tubes of equal cells for DNA and RNA extractions.

To check for deletions, we used the following pairs of primers: RET-7 (*RET-7_FWD 5’ GAGGAACCGCGCACATCAG 3’* and *RET-7_REV 5’ CGCCTTGCCTGGCCCTCCA 3’*), RET-5.5 (*RET-5*.*5_FWD 5’ GTCCTGGTTGCCCCACTCTG 3’* and *RET-5*.*5_REV 5’ GGTCAGGTGTCACAAAGTCT 3’*) and RET+3 (*RET+3_FWD 5’ AATGGGCAAGACCATCTCAG 3*’ and RET+3_*REV 5’ GTGGCCAGTAGCTGGAAGAGCAG 3*’). The primers amplify a wildtype sequence of 1.1kb for each enhancer region. Each cell line was PCR confirmed for enhancer deletion and Sanger sequenced to re-confirm the precise region of deletion. For quantifying gene expression changes, RNA was extracted from each cell line with deletions and qPCR assays performed in triplicate; n=6 for RET-7 and n=9 for RET-5.5 and RET+3. Quantitative gene expression changes were calculated as detailed above.

### Estimating haplotype-specific HSCR risk

Genotypes at 10 RET CRE variants in 220 S-HSCR cases and 503 European-ancestry controls were obtained from our published HSCR Genome Wide Association study ^14^ and the 1000 Genomes project ^21^, respectively. Haplotypes were generated from unphased genotypes using BEAGLE ^35^ and filtered to retain only those that had a frequency >1% in controls. Standard methods using χ^2^ statistics were used to calculate haplotype-count based odds ratios (OR), their upper and lower confidence limits and significance of their deviation from the null hypothesis of no association (OR = 1) ^13^.

### Identifying transcription factors for candidate enhancers

We searched for TF binding sites (TFBS) within all putative CREs using FIMO ^36,37^ and 890 validated TF motifs in TRANSFAC ^38^. We used the setting of “minimize false positives” and a stringent cut off of P<10^−4^ to identify candidate cognate TFs.

## Author Contributions

S.C and A.C. conceived and designed the study. K.M.K. conducted all *in vitro* luciferase assays, A.K. all genotyping assays and S.C and L.E.F all *in vivo* CRISPR assays. S.C. and A.C. wrote and edited the manuscript.

