## Supplemental Tables 1-3 for "A multi-enhancer *RET* regulatory code is disrupted in Hirschsprung disease"

| Enhancer | Percentage of cells with no deletion | Percentage of cells with 1-3 bp deletion | Percentage of cells with >3 bp deletion |
|---|---|---|---|
| E2 | 30 | 52 | 18 |
| E4 | 68 | 22 | 10 |
| E5 | 100 | 0 | 0 |
| E14 | 69 | 20 | 11 |
| E26 | 42 | 24 | 34 |
| E27 | 72 | 15 | 13 |
| E28 | 63 | 13 | 24 |
| RET-7 | 80 | 5 | 15 |
| RET-5.5 | 60 | 7 | 33 |
| RET+3 | 61 | 20 | 19 |

**Supplementary Table 1:** Percentage of cells carrying deletions of various length calculated using *Inference of CRISPR Edits (ICE)* tool (Hsiau et al., 2019). Nucleotides surrounding the HSCR associated polymorphisms in all enhancers were successfully deleted except for enhancer 5.

| Enhancer | Guide RNA Sequence |
|---|---|
| E2 | TAAACCGTTAAGTAATGACC |
| E4 | TGGGACTCATCTGTGCACGG |
| E5 | AGTGAGCCAGCAAAGTGAAA |
| E14 | CAGCCCGCCTCACTGCTCCA |
| E26 | GTGCTGAGCAAACAAGCCTG |
| E27 | GGCACCCTGTAAGAGTGAGG |
| E28 | AGCCCCTGCCTCCACAGACA |
| RET-7 | AGAAACCAGCAGAGCCACAT |
| RET-5.5 | GTGAGAAGCTGAAAGAGTGT |
| RET+3 | ACCCTTACATGGTCATCCAC |

**Supplementary Table 2:** Sequences of the individual guide RNA targeting the HSCR associated polymorphisms in the 10 enhancers.

| Enhancer | Forward Primer | Reverse Primer | Amplicon Size |
|---|---|---|---|
| E2 | CTTCAAACCGCACCCGCTCCTT | AGTGACGTCCTGCCTGCATCCT | 283 |
| E4 | AGCCTGAATCCAGCCACCAGGT | CTGCCCTGGGAGAGTGTCCACA | 417 |
| E5 | CAGCCAGCCCTCAGCACATGTC | AGAGTGACCTGCCAGCTCCCTG | 525 |
| E14 | AGCAGGCCTCTTGCTCAGAGCT | AGTGGGAGAGGCAGTCAGCTGG | 538 |
| E26 | TGCAGGTGGTCCTACCCGAGTG | GGAGTGTGAAAGGGCTGGGCAC | 339 |
| E27 | GTGCCCAGCCCTTTCACACTCC | CCCCACCTCACTGCTCAGGTCA | 421 |
| E28 | TCGCATGAGGGGTTCTGCAGGA | CCATTACACACCACGGGGGCAC | 512 |
| RET-7 | CCCCTGCTCCGTGGAACCCTAA | CCTGGCAGTGCTCAACAGACGG | 455 |
| RET-5.5 | AGCCTGTGATGCTGGGCTCTCA | GCCAGAGCTGAGCTGGGAGACT | 349 |
| RET+3 | CTGGGGTAGGGCGAAAATGGGC | ACCCAGCCTTCTGCTCTGAGGG | 537 |

**Supplementary Table 3:** PCR primer sequences used for Sanger sequencing to detect CRISPR induced insertions and deletions surrounding the HSCR associated polymorphisms in the 10 enhancers.
